# Rapid dexamethasone treatment inhibits LPS-induced cytokine storm in mice

**DOI:** 10.1101/2023.12.07.570372

**Authors:** Fen Zhang, Lanlan Xiao, Yujia Li, Rilu Feng, Menghao Zhou, Shima Tang, Roman Liebe, Matthias P Ebert, Steven Dooley, Lanjuan Li, Hong-Lei Weng

## Abstract

Severe infection-induced cytokine storm is an urgent medical syndrome with high mortality. To date, no therapy is available. This study shows that high concentrations of lipopolysaccharide (LPS) induce cytokine storm within 48h and thus kill most experimental mice. Rapid, but not late dexamethasone administration remarkably inhibits cytokine storm and rescues LPS-treated mice. Monocytes and macrophages are the major source of cytokine storm. In these cells, pro-inflammatory genes (i.e., *Tnf*, *Il6* and *Il1β*) have preassembled RNA polymerase II (RNA Pol II), but stay at the pause stage of transcriptional elongation in the absence of stimulation. LPS rapidly activates transcription of these “pre-loaded” genes within 2h. Administration of dexamethasone within this time window inhibits RNA Pol II ser2 binding to the core promoters of pro-inflammatory genes and thus reduces LPS-induced cytokine transcription. Therefore, rapid utilization of dexamethasone might be efficacious to prevent severe bacterium-induced cytokine storm in clinical practice.

## Introduction

Cytokine storm is a life-threaten systemic inflammatory syndrome that results in epithelial damage, capillary leakage, and multi-organ injury [1]. It can be triggered by a variety of pathogens such as bacterial and viral infections, immunotherapy, and autoimmune diseases [2]. In the condition of bacterial infection-induced cytokine storm, the most prominent cytokines are IL-6, IL-1, and TNF-α. They are mainly secreted by macrophages and monocytes and elicit fever through distinct mechanisms [3]. High levels of TNF-α are lethal to the host organism [4].

The detailed mechanisms of cytokine induction in the condition of bacterial infection remain largely unknown to date. Macrophages play a leading role in the occurrence of cytokine storm [5]. Cytokine expression in macrophages is orchestrated by transcription factors. Medzhitov divided transcription factors in macrophages into three classes: Class I, constitutively expressed but latent; Class II, synthesized de novo; and Class III, induced during cell differentiation [6]. These three categories of transcription factors control the successive expression of primary and secondary response genes when macrophages are stimulated by bacterial components such as lipopolysaccharide (LPS). Pro-inflammatory cytokine genes are governed by Class I transcription factors, in particular, nuclear factor-κB (NF-κB) and interferon-regulatory factor (IRF) [6]. It is noteworthy that the transcription of primary response genes is completed in the period between 0.5h and 2h following LPS stimulation [6]. These observations imply that the occurrence of cytokine storm might be quite rapid when the host is challenged by severe bacterial infection.

The transcription of inflammatory genes is negatively regulated by multiple factors, including nuclear receptors such as glucocorticoid receptors. The activation of the glucocorticoid receptor increases macrophage uptake of apoptotic cells while concurrently inhibiting inflammatory signaling pathways [7]. In addition, activated glucocorticoid receptor inhibits chromatin occupancy of NF-κB and thus influences LPS-induced gene transcription in macrophages [8]. These studies suggest that glucocorticoid might be a potential candidate to inhibit cytokine storm.

In this study, investigation focuses on the effect of dexamethasone (DEX) [9] on the regulation of cytokine storm in mice treated with high concentrations of LPS.

## Material and methods

### Reagents

The primers, antibodies, and chemical reagents used in this study are described in the **Table 1-3**.

**Table 1.**
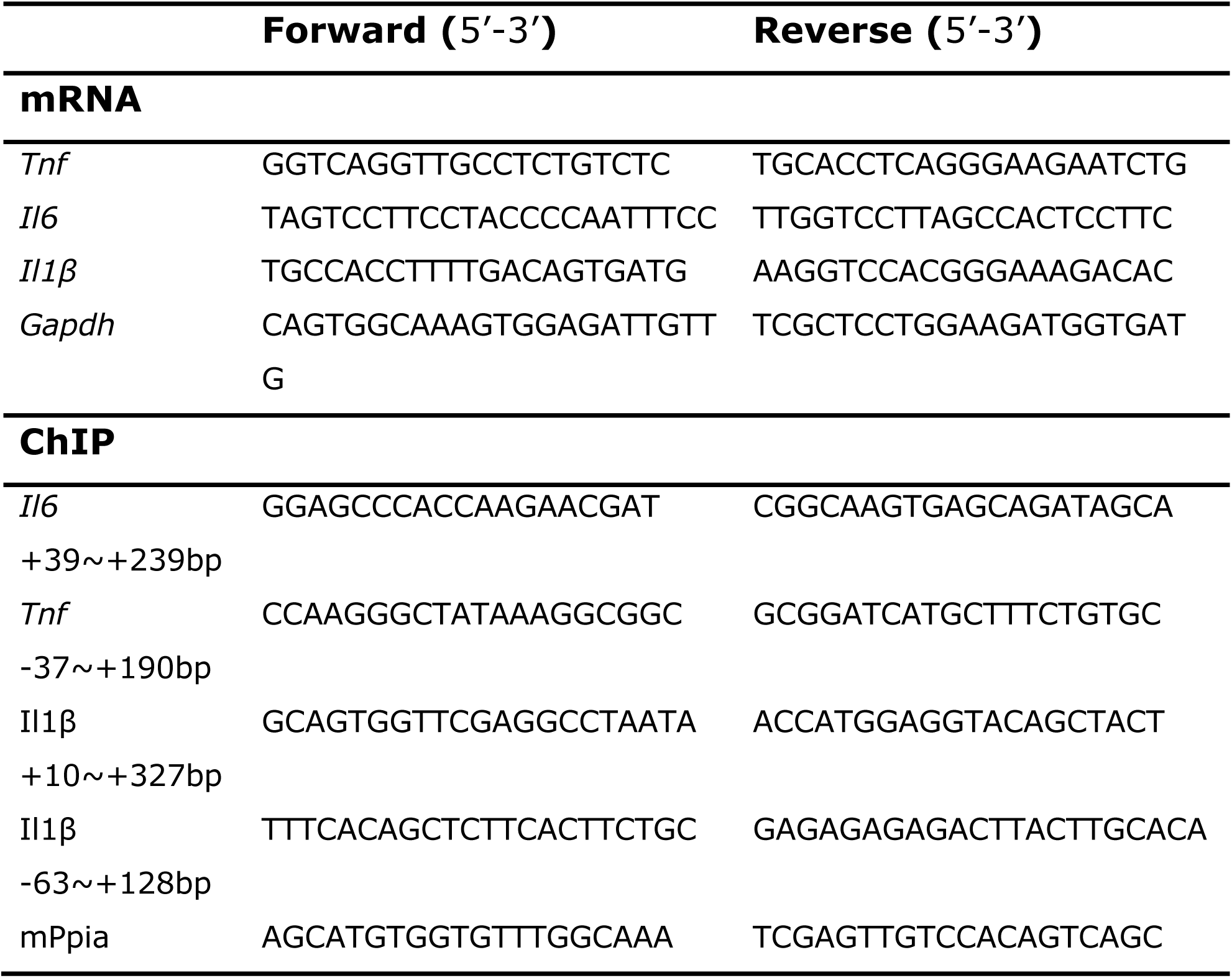
Primer sequences.

**Table 2.** Antibodies Used in the Study.

| **ChIP assay** |  |  |  |
| --- | --- | --- | --- |
| RNA PolII Ser2 | Rabbit | Abcam | Ab193468 |
| RNA PolII Ser5 | Rabbit | Abcam | Ab5131 |
| IgG Control | Rabbit | Cell Signaling Technology | CST 3900 |
| ChIP buffer | 1 mM EDTA at pH 8.0, 50 mM HEPES-KOH at pH 7.5, 1x protease inhibitors, 140 mM NaCl, 1% Triton X-100, 0.1% sodium deoxycholate, 0.1% SDS |  |  |
| RIPA buffer | 2 mM EDTA at pH 8.0, 20 mM Tris-HCl at pH 8.0, 1% Triton X-100, 150 mM NaCl, 1x protease inhibitors |  |  |
| Low salt wash buffer | 2mM EDTA at pH 8.0, 20mM Tris-HCl at pH 8.0, 150mM NaCl, 1% Triton X-100, 0.1% SDS |  |  |
| High salt wash buffer | 2mM EDTA at pH 8.0, 20mM Tris-HCl at pH 8.0, 500mM NaCl, 1% Triton X-100, 0.1% SDS |  |  |
| LiCl wash buffer | 1 mM EDTA at pH 8.0, 10mM Tris-HCl at pH 8.0, 0.25M LiCl, 1% NP-40, 1% sodium deoxycholate |  |  |
| TE buffer | 1mM EDTA at pH 8.0, 10mM Tris-HCl at pH 8.0 |  |  |

**Table 3.**
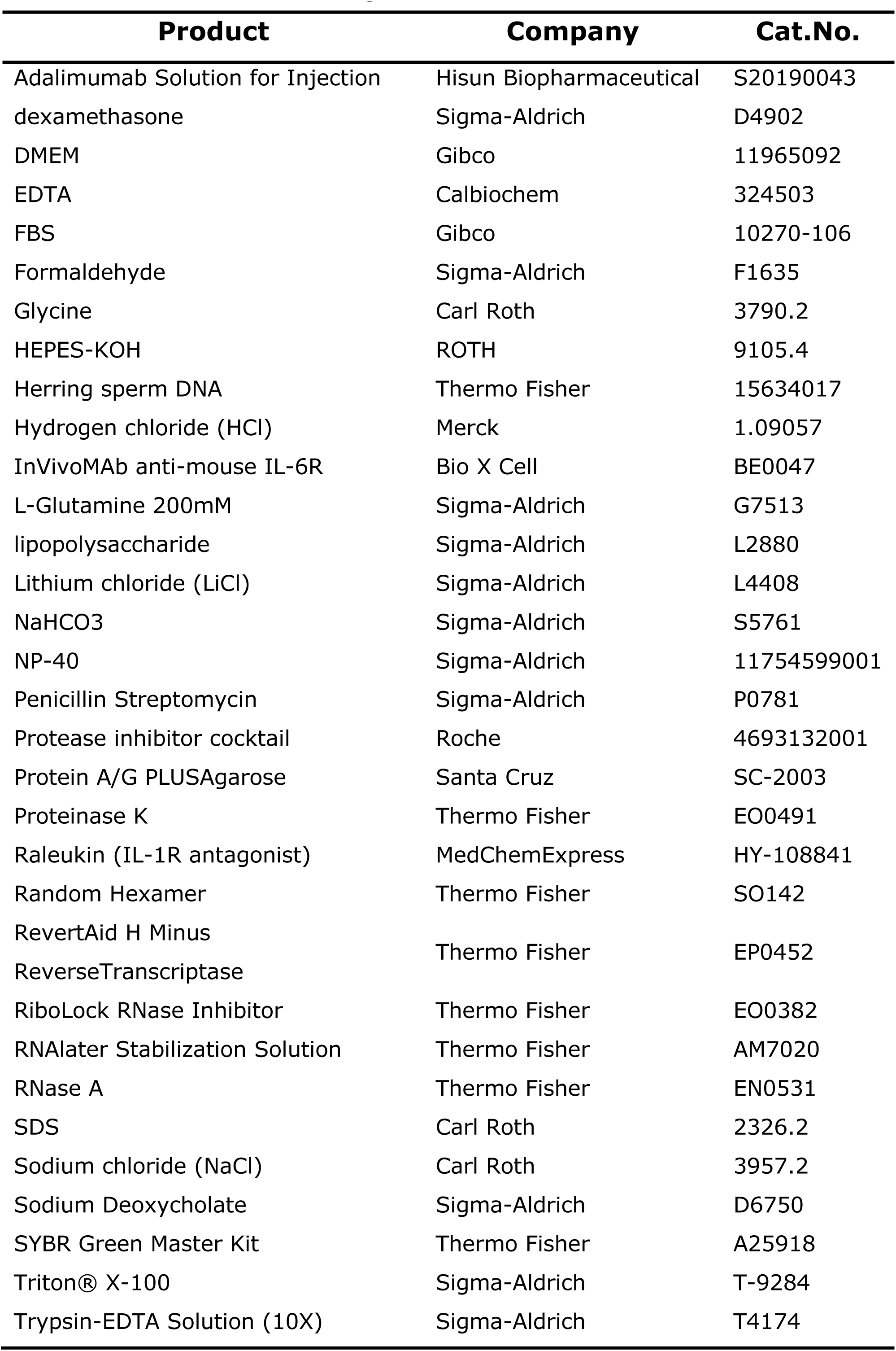
Chemicals and reagents.

### Animals

The animal experiment was approved by the Animal Care Committee of the Animal Experimental Ethical Inspection of the First Affiliated Hospital, College of Medicine, Zhejiang University (No. 2022-1618). 156 male, 4-6 week old C57BL/6 mice were sourced from the laboratory at the Animal Center of the College of Medicine, First Affiliated Hospital, Zhejiang University. Food and water were provided ad libitum with a standard condition of 12-hour photoperiod.

After acclimating in the laboratory animal center over one week, mice were administered intraperitoneal injections of PBS, 5 mg/kg DEX, LPS (30 mg/kg, Sigma-Aldrich, Darmstadt, Germany), or a combination of LPS and DEX, as depicted in Figure 1A. After the experimental period, mice were humanely euthanized under isoflurane anesthesia. Heart, lung, liver, kidney, and serum samples were collected.

**Fig. 1:**
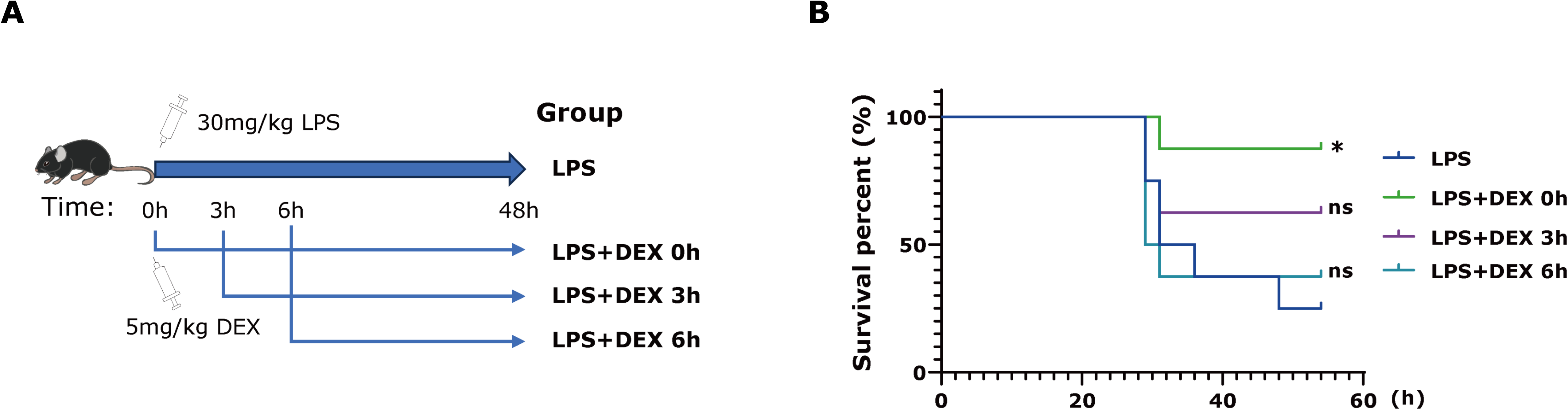
Rapid administration of dexamethasone rescues LPS-induced mouse death. (**A**) A scheme depicts administration of LPS and dexamethasone in mice. (**B**) Kaplan-Meier plot shows survival rates of mice with indicated treatment (n=8 per group). ns, no significance; *, *P*<0.05; **, *P*<0.01; ***, *P*<0.001 compared with LPS group.

### Cells

The Raw264.7 mouse macrophage cell line was used in our vitro experiment and cultured in DMEM medium supplemented with 10% FBS (Gibco, New York, USA), 1% L-glutamine, and 100 U/ml penicillin-streptomycin. Raw264.7 cells were treated with LPS (200ng/ml) and DEX (6µM).

### Hematoxylin and eosin staining

Tissues were collected and fixed with 4% paraformaldehyde overnight before being embedded in paraffin. Slices with a thickness of 3-5µm were utilized for the staining of hematoxylin and eosin (H&E). The resulting sections were scanned using NanoZoomer 2.0-RS by HAMAMATSU PHOTONICS K.K. in Tokyo, Japan.

### Enzyme-linked immunosorbent assay

The serum TNF-α, IL-6, and IL-1β concentrations were determined using commercial Enzyme-linked immunosorbent assay (ELISA) kits (Beijing Dakwei Biotechnology, Beijing, China) according to the vendor’s instructions. OD450 readings were obtained with a microplate reader (Bio-Rad, California, USA).

### RNA sequencing and bioinformatics analysis

RNA sequencing (RNA-seq) of peripheral blood mononuclear cells (PBMCs) was performed based on the SMART-Seq protocol. Briefly, RNA was extracted from PBMCs and reverse transcribed to cDNA. Modified oligo(dT) primer (the SMART CDS Primer) primed the first-strand synthesis reaction. Subsequently, single strand cDNA was amplified with LD PCR to generate sufficient dscDNA for library construction. cDNA was fragmented with dsDNA Fragmentase (NEB, M0348S, Massachusetts, USA) for 30 minutes at 37℃. The fragmented cDNA was used to construct the NGS library. Blunt-ended DNA fragments were produced by combining fill-in reactions with exonuclease activity. Sample purification beads were used to perform size selection. A nucleotide was added and then indexed Y adapters were ligated to the fragments in order to be amplified through PCR. The paired-end sequencing was conducted with Illumina Novaseq 6000 (Illumina, California, USA), following the recommended protocol from the vendor to create raw data.

The filtered raw data were compared to the reference sequence to determine gene expression levels. Based on gene expression in different sample groups, differentially expressed genes (DEG) with *p*<0.05 were identified. The volcano, venn and KEGG enrichment analysis were calculated based on the R (https://www.r-project.org/) on the OmicStudio platform (https://www.omicstudio.cn/tool) [10].

### RNA extraction and quantitative real-time PCR

Total RNA was extracted with TRIzol (Invitrogen Life Technologies, Massachusetts, USA). RNA concentrations were detected by NanoDrop (Thermo Fisher Scientific, Massachusetts, USA). Reverse transcription to synthesize cDNA was performed using RevertAid H Minus Reverse Transcriptase. The quantitative real-time PCR (qRT-PCR) experiments were performed utilizing the POWRUP SYBR MASTER MIX via a StepOnePlus Real-time PCR instrument (Applied Biosystems, Texas, USA).

### Chromatin immunoprecipitation and quantitative real-time PCR

Raw264.7 cells were cross-linked with 1% formaldehyde for 10 minutes and followed by 125 mM glycine treatment at room temperature for 5 minutes. The cells were washed twice using cold PBS, collected and transferred to tubes, centrifuged at 4℃, and then resuspended in ChIP buffer. Incubation was carried out on ice for 10 minutes. The cell pellet was fragmented by sonicator (Diagenode, New Zealand, USA) using 30-second on/off cycles for 30 repetitions, resulting in an average fragment size of 500 base pairs of DNA. The supernatant was collected through centrifugation at 8000g for 10 minutes at 4℃. 50µL of each fragmented sample was utilized as input. Immunoprecipitation samples were diluted with RIPA buffer and then incubated overnight at 4℃ with 5µg of primary antibody or IgG on a rotating device. 60µL of Protein A/G PLUS-agarose beads (Santa Cruz Biotechnology, Texas, USA) and 4µg of single-stranded herring sperm DNA were added to each sample. The samples were then incubated on a rotating device at 4℃ for 3-4 hours. After centrifugation, the supernatant was removed, and the beads were washed for 10 minutes at 4℃ with following buffers sequentially: low salt wash buffer, high salt wash buffer, LiCl wash buffer, and TE buffer. Subsequently, the beads were suspended in 120µL of elution buffer, freshly prepared with 1% SDS and 100mM NaHCO3, and gently agitated for 15 minutes at a temperature of 30℃. After centrifugation, the supernatant and input samples underwent incubation with 4.8µL of 5M NaCl and 2µL of 10mg/mL RNase A at 65℃ overnight to reverse the crosslinks. Then, the samples were incubated with 2µL of proteinase K (20mg/mL) at 60℃ for 1 hour. All samples underwent purification with a MinElute PCR Purification Kit (Qiagen, DE) and subsequent analysis with POWRUP SYBR MASTER MIX using a StepOnePlus Real-time PCR instrument.

### Antagonists for TNF-α, IL-6, and IL-1β

To inhibit individual cytokine in LPS-treated mice, Adalimumab solution for injection (anti-TNF-α, 10mg/kg dosage, Hisun Biopharmaceutical, Taizhou, China) [11], InVivoMAb anti-mouse IL-6R (8mg/kg dosage, Bio X Cell, New Hampshire, USA) [12], and Raleukin (IL-1R antagonist, 10 or 20mg/kg dosage, MedChemExpress, New Jersey, USA) [13] were used.

### Statistical analysis

Two-tailed unpaired t-test was employed to compare differences between two groups, whereas one-way ANOVA was used for more than two groups. Log-rank (Mantel-Cox) test was utilized to compare the survival rates between two groups. Statistical analyses and figures were performed using Prism version 9.3.1 (GraphPad Software, San Diego, USA). Continuous variables are presented as mean ± standard deviation (mean ± SD). Statistical significance was set at *P*<0.05 and is denoted in the figures with *, *P*<0.05; **, *P*<0.01; ***, *P*<0.001.

## Results

### Rapid administration of dexamethasone rescues LPS-treated mice

To examine the effects of DEX on LPS treatment, mice were injected with high concentrations of LPS (30mg/kg body weight, **Figure 1A**). Such a high dose of LPS killed 6 out of 8 mice (75%) within 48h (**Figure 1B**). Impressively, injection of 5mg/kg DEX simultaneously with LPS reduced mouse mortality to 12.5% (**Figure 1B**). The mortality of mice increased to 37.5% and 62.5% respectively when mice received DEX 3h and 6h following LPS administration (**Figure 1B**).

These results suggest that rapid utilization of DEX is critical for rescuing mice from LPS-induced death.

### Rapid dexamethasone treatment inhibits cytokine storm

Subsequently, pathological changes of vital organs were scrutinized in LPS-treated mice. Histological examination did not find severe damages in the heart, lung, liver, and kidney collected from the dead mice (**Figure 2**). Serum ELISA assays showed that LPS administration rapidly induced high levels of cytokines, including IL-6, TNF-α and IL-1β (**Figure 3A-C**). The peak levels of IL-6 and TNF-α were at 3h following LPS treatment, while the peak level of IL-1β was at 12h. The average concentrations in the peak levels of IL-6, TNF-α and IL-1β were 646.6±194.3ng/mL, 1546±472.7ng/mL and 14.95±4.123pg/mL, respectively (**Figure 3A-C**). Afterwards, the levels of both cytokines were reduced rapidly. The concentrations of IL-6 and TNF-α reduced to 234.9±142ng/mL, 61±7.788ng/mL, and 32.39±7.833ng/mL and 365.2±97.43ng/mL, 170.5±20.88ng/mL, and 369.7±142.1ng/mL, at 6h, 12h and 18h following LPS treatment.

**Fig. 2:**
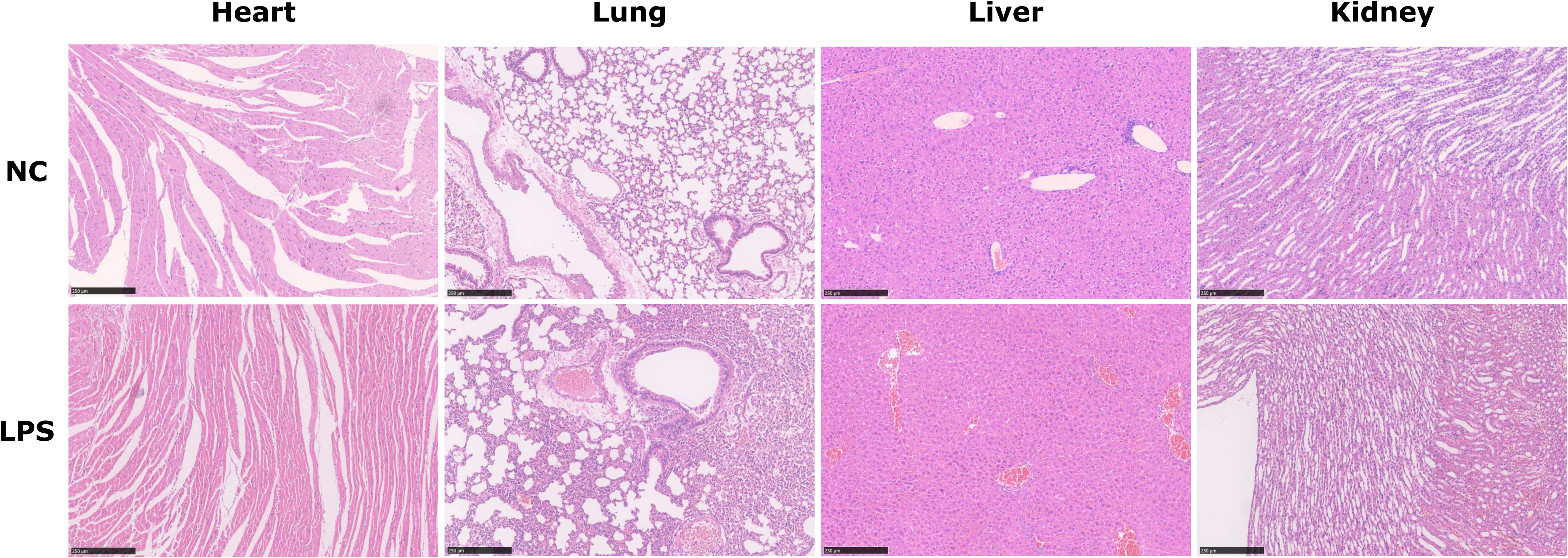
Pathological changes of vital organs in LPS-treated mice and control mice. H&E staining was performed in mice with or without LPS injection. Representative images show histological alterations in the heart, lung, liver and kidney. Scale bars: 250µm.

**Fig. 3:**
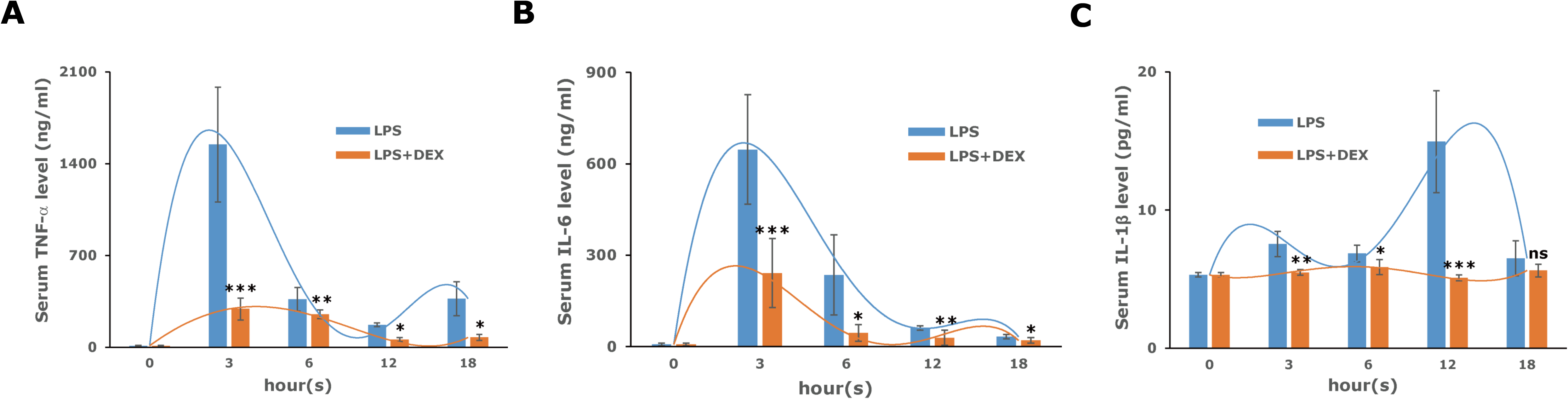
Dexamethasone reduces LPS-induced blood pro-inflammatory cytokine levels in mice. Serum (**A**) TNF-α, (**B**) IL-6 and (**C**) IL-1β levels were examined in mice with indicated treatment. Bars represent the mean ± SD. *P*-values were calculated by unpaired Student’s t test, ns, no significance; *, *P*<0.05; **, *P*<0.01; ***, *P*<0.001 compared with LPS group.

DEX remarkably reduced IL-6 concentrations to 241.3±122.4ng/mL, 44.62±29.86ng/mL, 29.14±27.18ng/mL and 19.96±10.23ng/mL at 3h, 6h, 12h and 18h, respectively and decreased TNF-α to 291.1±89.47ng/mL, 251.1±34.55ng/mL, 56.78±18.68ng/mL and 74.33±28.04ng/mL at 3h, 6h, 12h and 18h (**Figure 3A-B**). In addition, DEX remarkably reduced IL-1β concentrations to 5.10±0.24pg/mL at 12h (**Figure 3C**).

There results suggest that high concentrations of LPS-induced lethal cytokine storm occurs very rapid. Administration of DEX only in early phase can inhibit cytokine storm.

### Alteration of inflammatory signaling pathways is prominent in mouse peripheral blood monocytes (PBMCs)

Given that monocytes are the main cell source producing pro-inflammatory cytokines, RNA-seq was subsequently performed in PBMCs collected from LPS-treated mice with or without DEX treatment. Following LPS + DEX treatment for 3h, 1306 transcripts, including pro-inflammatory cytokine IL-1β, were altered in PBMCs compared with those collected from LPS-treated mice (**Figure 4A**). KEGG enrichment analyses showed 10 top signaling pathways significantly influenced by DEX in LPS-treated mice (**Figure 4B**). Following LPS treatment for 3h, 6660 transcripts were altered in PBMCs. DEX administration influenced 1306 transcripts, including 708 LPS-impacted genes (**Figure 4C**). KEGG enrichment analyses further showed 10 signaling pathways remarkably influenced by the 708 LPS and DEX impacted genes (**Figure 4D**). Among them, TNF-α, NOD-like receptor and Toll-like receptor signaling pathways were significantly altered by DEX administration (**Figure 4B and 4D**).

**Fig. 4:**
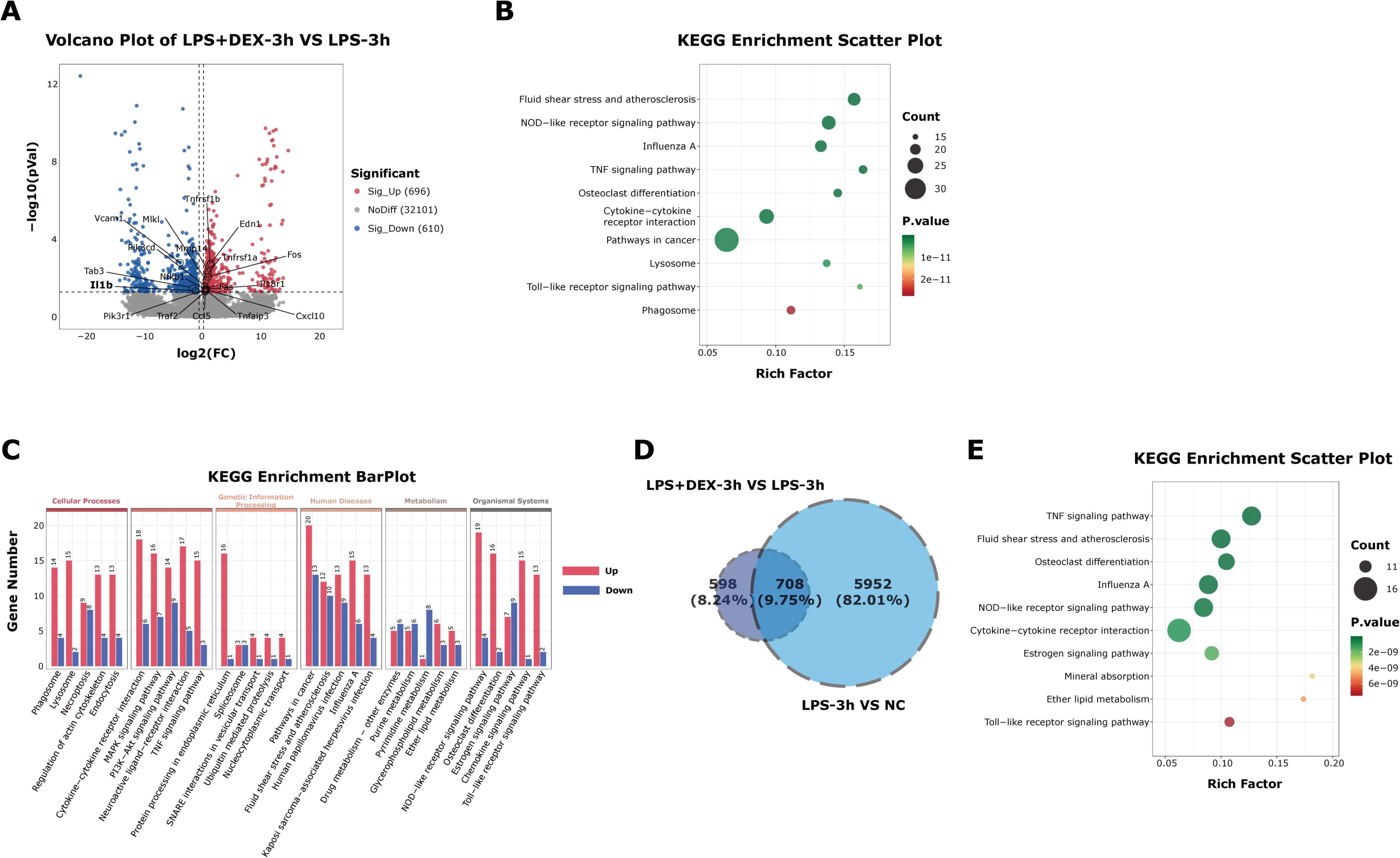
Transcriptome alterations in peripheral blood monocytes (PBMCs) from LPS-treated mice with or without dexamethasone administration. RNA-seq was performed in PBMCs collected from mice with different treatments (n=3 each group). (**A**) Volcano plot shows differentially expressed RNAs between LPS-treated mice with or without DEX for 3h. (**B**) KEGG enrichment analyzed enriched pathways in PBMCs collected from LPS-treated mice with or without DEX for 3h. (**C**) Venn diagram depicts differentially expressed genes between normal mice, mice treated with LPS for 3h, and mice simultaneously treated with LPS and DEX for 3h. 708 overlapped genes were identified. (**D**) KEGG enrichment analyzed pathways the 708 overlapped genes influenced.

### Dexamethasone-inhibited cytokine expression is time-dependent in vitro

To further clarify the effects of DEX on cytokine expression in macrophages, expression of *Tnf*, *Il6* and *Il1β* was examined in mouse macrophage line RAW264.7 cells. Consistent with the in vivo observations, administration of DEX at different time points yielded different outcomes. Administration of DEX simultaneously with LPS or 0.5h following LPS injection reduced mRNA expression of *Tnf*, *Il6* and *Il1β* more efficiently than administration at 2.5h after LPS injection (**Figure 5A-C**). The administration of DEX simultaneously with LPS demonstrated the highest efficiency at inhibiting expression of these pro-inflammatory cytokines.

**Fig. 5:**
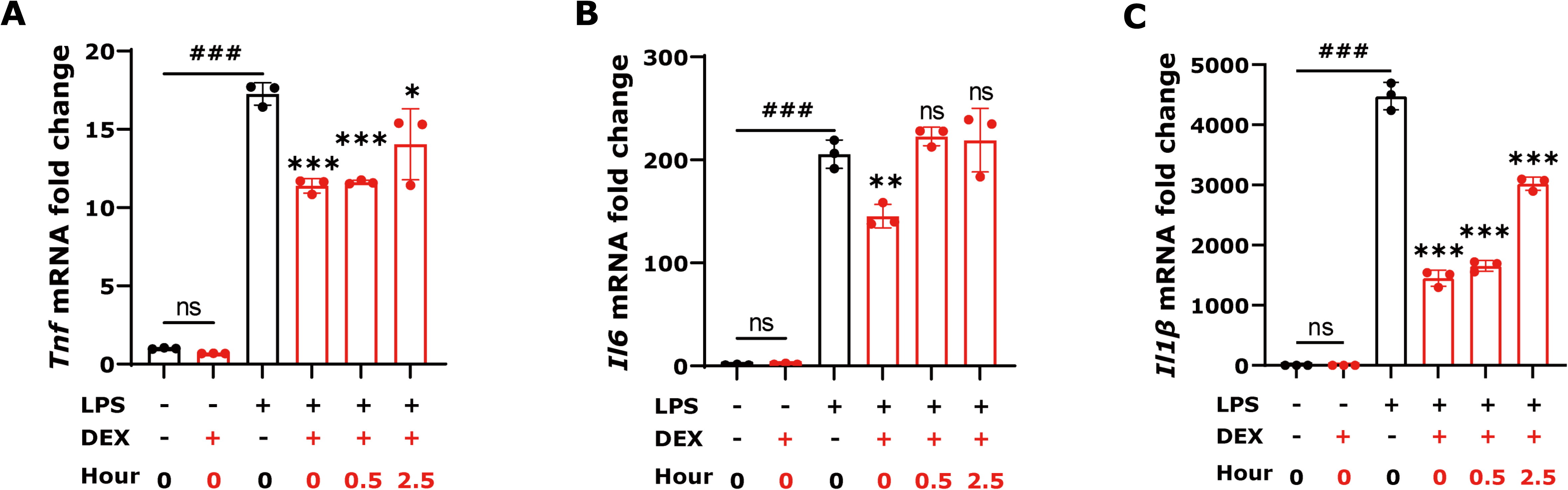
Dexamethasone reduces LPS-induced pro-inflammatory cytokine mRNA expression in RAW264.7 cells. qPCR was performed to examine (**A**) *Tnfa*, (**B**) *Il6* and (**C**) *Il1β* mRNA expression in RAW264.7 cells with indicated treatment. The “hour(s)” indicates the time that DEX added simultaneously with LPS (0), or after LPS treatment (0.5 and 2.5). Bars represent the mean ± SD. *P*-values were calculated by unpaired Student’s t test, ns, no significance; *, *P*<0.05; **, *P*<0.01; ***, *P*<0.001 compared with LPS group. ###, *P*<0.001 compared with control group.

### Rapid dexamethasone administration inhibits transcription elongation of pro-inflammatory cytokine genes through reducing RNA Pol II ser2

To clarify how rapid DEX administration is capable of inhibiting pro-inflammatory cytokine gene expression, ChIP assays were performed to observe the binding of RNA Pol II to phosphorylated serine 5 or serine 2 of the heptapeptide in the C-terminal domain (RNA Pol II ser5, **Figure 6A-C** or RNA Pol II ser2, **Figure 6D-E**) of the *Tnfa*, *Il6* and *Il1β* promoters in RAW264.7 cells with LPS and DEX treatment. RNA Pol II ser5 and RNA Pol II ser2 occurs in transcription initiation and elongation phases, respectively [14, 15]. In RAW264.7 cells simultaneously treated with LPS and DEX for 3h, LPS-induced RNA Pol II ser5 binding to the promoters of *Tnfa* and *Il1β* (**Figure 6A and 6C**), and RNA Pol II ser2 binding to the promoters of *Tnfa*, *Il6* and *Il1β* were significantly reduced (**Figure 6D-E**). In contrast to the administration of DEX with LPS simultaneously, DEX treatment 2.5h following LPS only reduced RNA Pol II ser5 binding to the promoters of *Tnfa* (**Figure 6A**) and was not capable of reducing RNA Pol II ser2 binding to the promoters of three pro-inflammatory genes (**Figure 6D**). These results suggest that 2h is a time threshold for DEX to inhibit LPS-induced pro-inflammatory gene transcription elongation.

**Fig. 6:**
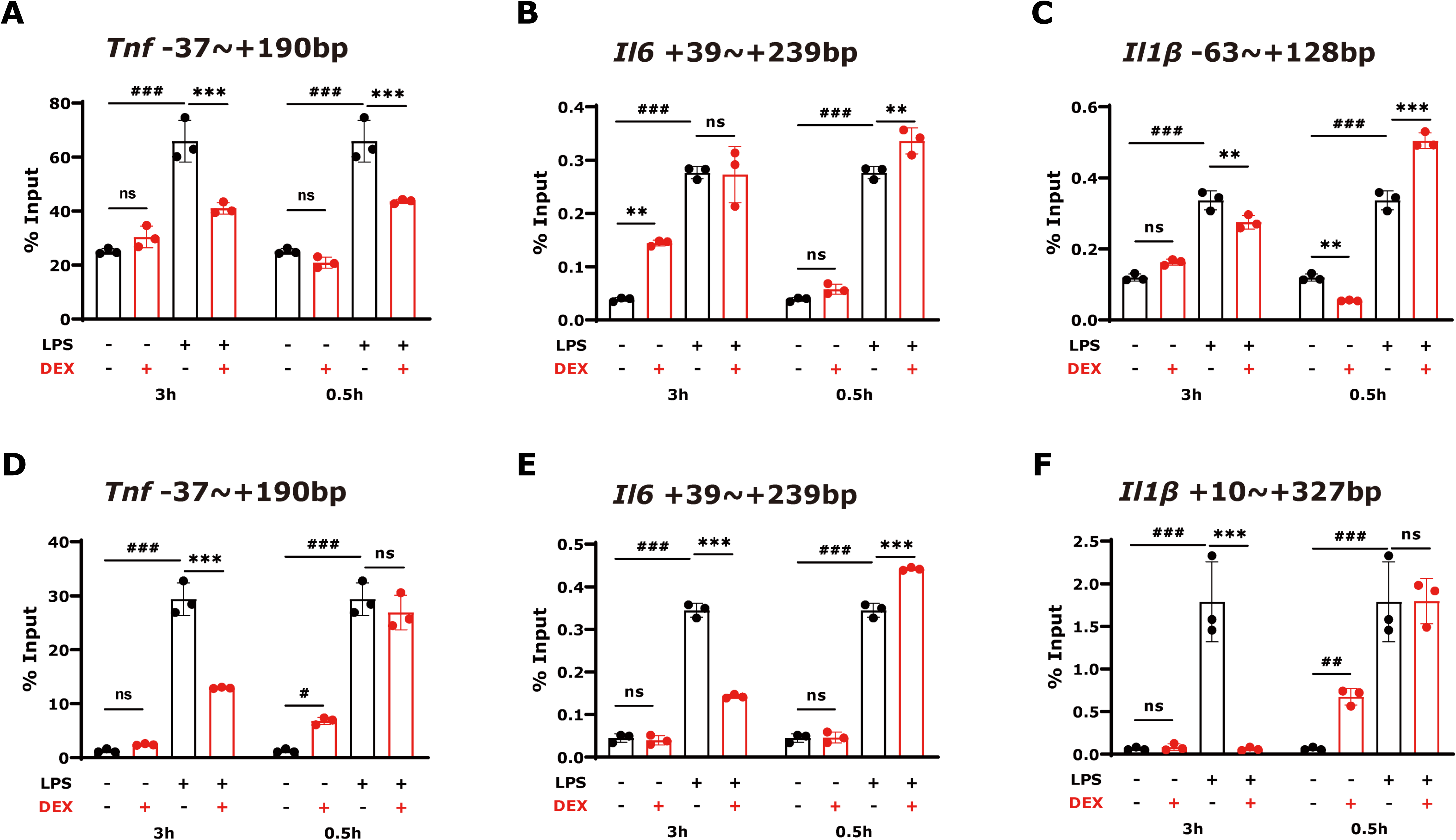
The effects of dexamethasone on the LPS-induced pro-inflammatory cytokine transcription in macrophages. Chromatin immunoprecipitation (ChIP) assays for RNA polymerase II phosphorylation of Ser5 (Pol II ser5, **A-C**) and Ser2 (Pol II ser2, **D-F**) were performed in RAW264.7 cells. In the left panels, RAW264.7 cells were simultaneously treated by LPS and dexamethasone for 3h. In the right panels, the cells were first treated by LPS for 2.5h. Subsequently, added dexamethasone to the cells for additional 0.5h. Bars represent the mean ± SD. *P*-values were calculated by unpaired Student’s t test. ns, no significance; *, *P*<0.05; **, *P*<0.01; ***, *P*<0.001 compared with LPS group. #, *P*<0.05; ##, *P*<0.01; ###, *P*<0.001 compared with control group.

### Inhibition of TNF-α, but not IL-6 and IL-1β, rescue mice from death

In clinical practice, clinicians usually do not have a chance to treat patients within 2h following severe bacterial infection. How to rescue these patients from lethal cytokine storm? One may speculate that inhibiting individual TNF-α, IL-6 or IL-1 can rescue patients. To answer this question, the impact of Adalimumab (**Figure 7A**), InVivoMAb anti-mouse IL-6R (**Figure 7B**) and Raleukin (**Figure 7C-D**) was analyzed. Only TNF-α antagonist Adalimumab was capable of improving LPS-induced mouse mortality (**Figure 7A**). These results suggest a critical role of TNF-α in cytokine storm-caused death.

**Fig. 7:**
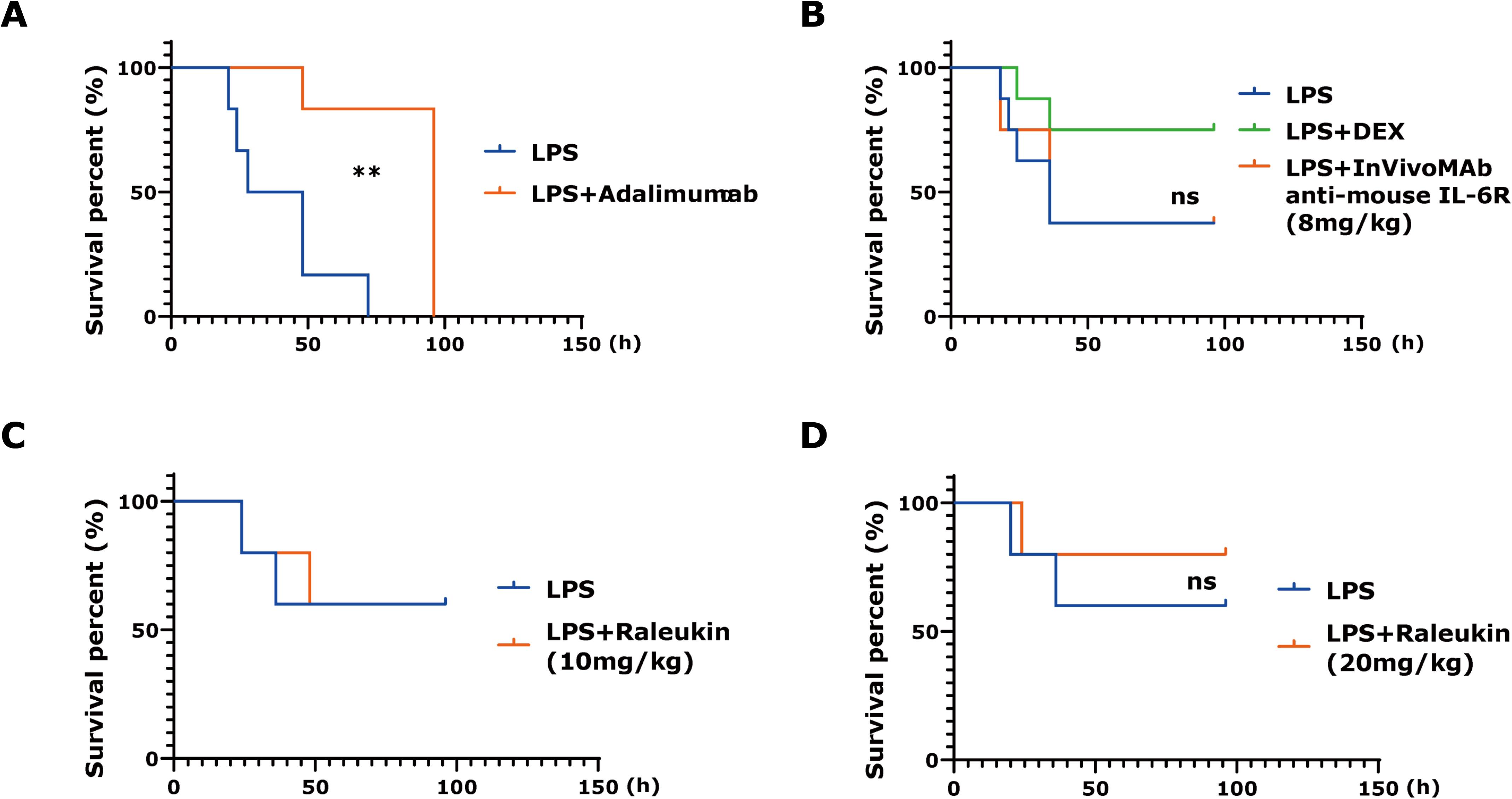
Inhibiting TNF-α, but not IL-6 and IL-1, can rescue mice from high concentrations of LPS-caused death. Kaplan-Meier plot shows survival rates of mice with indicated treatment. (**A**) Adalimumab Solution for Injection (anti-TNF-α, 10mg/kg dosage, n=6 per group), (**B**) InVivoMAb anti-mouse IL-6R (8mg/kg dosage, n=6 in LPS and LPS+DEX groups and n=8 in LPS+anti-IL-6R group), and (**C-D**) Raleukin (IL-1R antagonist, 10 or 20mg/kg dosage, n=5 per group) were used in LPS mice. ns, no significance; *, *P*<0.05; **, *P*<0.01; ***, *P*<0.001 compared with LPS group.

## Discussion

In severe bacterial infection, successful inflammatory response is of vital importance to eliminate invading bacteria. The magnitude and duration of the inflammatory response depend on the bacterial load. Excessive inflammatory response such as cytokine storm is essential to clear pathogens, however, such a heavy response may result in severe and potentially lethal collateral damage [16]. When bacterial infection occurs, macrophages and peripheral monocytes are the major cellular sources of cytokines [6, 17–19]. It has been elegantly shown by Hargreaves et al. that many pro-inflammatory genes in macrophages are primary response genes, which have preassembled RNA Pol II at their promoters in the basal state [20]. In the absence of stimulation, RNA Pol II only generates low-level of full-length unspliced transcripts. In the condition of gram-negative bacterial infection, LPS rapidly initiates transcription elongation to generate mature and protein-coding transcripts through NFκB-dependent recruitment of TrxG protein bromodomain-containing 4 (Brd4) and P-TEFb, a complex composed of the kinase cdk9 and cyclin T in macrophages [20]. Therefore, pro-inflammatory cytokines in macrophages are rapidly produced upon LPS stimulation. Medzhitov divided the LPS-induced inflammatory response in macrophages into three phases characterized by three gene categories: primary response genes, secondary response genes and macrophage-specific genes [6, 21]. The primary response gene transcription occurs between 0.5h and 2h following LPS stimulation [6]. Most pro-inflammatory genes belong to the primary response genes. This explains why cytokine storm following severe bacterial infection occurs so fast. Consistent with these investigations, this study did observe extremely high levels of TNF-α (∼ 1500ng/mL) and IL-6 (∼ 600ng/mL) in PBMCs of mice treated with high concentrations of LPS for 3h. Subsequently, cytokine levels in PBMCs decline rapidly. It is not surprising that the majority of mice with excessive levels of TNF-α died within 48h following LPS injection [22].

To date, there is no commonly recognized therapy to treat cytokine storm [1, 23]. In clinical practice, it is controversial to adopt glucocorticoids to treat cytokine storm [8, 24]. Given a critical role of glucocorticoid receptor in the inhibition of chromatin occupancy of NF-κB in LPS-treated macrophages [8], the current study examined the effects of dexamethasone on mice treated with high concentrations of LPS. The most impressive observation is that only rapid utilization of DEX is capable of inhibiting cytokine storm and thus rescuing mice. The therapeutic efficiency is completely time-dependent. Subsequent mechanistic investigation showed that DEX cannot inhibit RNA Pol II ser2 binding to the promoters of TNF-α, IL-6 and IL-1β if macrophages were treated later than 2h after LPS stimulation. These observations suggest an essential effect of adopting glucocorticoid as early as possible to treat patients with severe bacterial infection in order to prevent cytokine storm-induced death.

In clinical practice, patients usually are not accessible for treatment during the golden time window 2h after bacterial infection [25]. After this window, glucocorticoid therapy is likely to fail. How to treat these patients? This study investigated the effects of individual cytokine inhibitors on LPS-treated mice. TNF-α, IL-6 and IL-1β are major components of cytokine storm induced by severe bacterial infection [1, 3, 23]. However, only inhibition of TNF-α shows potential therapeutic effect in the reduction of mortality of LPS-treated mice. Nosocomial infection is one of the leading causes resulting in sepsis and septic shock [26, 27]. For a hospitalized patient with a potential risk of secondary bacterial infection, whether clinicians should adopt glucocorticoids for prophylactic treatment is an important and interesting question.

Taken together, utilization of DEX in the early phase of LPS-treatment of mice significantly inhibits transcription elongation of pro-inflammatory genes in macrophages and thus reduces cytokine storm to rescue mice from death. These results strongly suggest that utilization of glucocorticoids within a limited time window might be essential to ameliorate severe bacterium-induced cytokine storm in clinical practice.

## Acknowledgements

We are grateful to Dr. Astrid Schmieder for the RAW 264.7 cell line.

## Abbreviations

ChIP: Chromatin immunoprecipitation
DEGs: differentially expressed genes
DEX: dexamethasone
DMEM: Dulbecco’s Modified Eagle Medium
ELISA: Enzyme-linked immunosorbent assay
FBS: Fatal bovine serum
H&E: Hematoxylin and eosin
LPS: lipopolysaccharide
NF-κB: nuclear factor-κB
IRF: interferon-regulatory factor
IL-1R: interleukin 1 receptor
IL-6R: interleukin 6 receptor
TNF-α: tumor necrosis factor alpha
FBS: fetal bovine serum
PBMC: peripheral blood mononuclear cell
qRT-PCR: quantitative real-time polymerase chain reaction
RNA-seq: RNA sequencing
SD: Standard deviation.

